# Non-severe burns induce a prolonged systemic metabolic phenotype indicative of a persistent inflammatory response post-injury

**DOI:** 10.1101/2023.04.24.537960

**Authors:** Monique J. Ryan, Edward Raby, Luke Whiley, Reika Masuda, Samantha Lodge, Philipp Nitschke, Garth L. Maker, Julien Wist, Elaine Holmes, Fiona M. Wood, Jeremy K. Nicholson, Mark W. Fear, Nicola Gray

## Abstract

Globally, burns are a significant cause of injury that can cause substantial acute trauma as well as lead to increased incidence of chronic co-morbidity and disease. To date, research has primarily focused on the systemic response to severe injury, with little in the literature reported on impact of non-severe injuries (<15% total burn surface area; TBSA). To elucidate the metabolic consequences of non-severe burn injury, longitudinal plasma was collected from adults (n=35) who presented at hospital with a non-severe burn injury at admission, and at 6 week follow up. A cross-sectional baseline sample was also collected from non-burn control participants (n=14). Samples underwent multiplatform metabolic phenotyping using ^1^H nuclear magnetic resonance spectroscopy and liquid chromatography-mass spectrometry to quantify 112 lipoprotein and glycoproteins signatures and 852 lipid species from across 20 subclasses.

Multivariate data modelling (Orthogonal projection to latent structures-discriminate analysis) revealed alterations in lipoprotein and lipid metabolism when comparing baseline control to hospital admission samples, with the phenotypic signature found to be sustained at follow up. Univariate (Mann-Whitney U) testing and OPLS-DA indicated specific increases in GlycB (p-value <1.0e^-4^), low density lipoprotein-2 subfractions (Variable importance in projection score; VIP >6.83e^-1^) and monoacyglyceride (20:4)(p-value <1.0e^-4^) and decreases in circulating anti-inflammatory high-density lipoprotein-4 subfractions (VIP >7.75e^-1^), phosphatidylcholines, phosphatidylglycerols, phosphatidylinositols and phosphatidylserines.

The results indicate a persistent systemic metabolic phenotype that occurs even in cases of non-severe burn injury. The phenotype is indicative of an acute inflammatory profile which continues to be sustained post-injury, suggesting an impact on systems health beyond the site of injury. The phenotypes contained metabolic signatures consistent with chronic inflammatory states reported to have elevated incidence post-burn injury. Such phenotypic signatures may provide patient stratification opportunities, to identify individual responses to injury, personalise intervention strategies and improve acute care, reducing risk of chronic co-morbidity.

## Introduction

Worldwide, burn injuries remain one of the most traumatic injuries for individuals to experience and, despite treatment advances, are associated with long-term impacts on physical and psychological health^1^. Thermal burn injury has been found to be associated with a plethora of long-term complications, including increased risk of cancer^2^, infectious^3^, cardiovascular disease^4^, diabetes mellitus^5^ and musculoskeletal disorders^6^. These systemic consequences of burn injury can occur even when there is little or no apparent scarring following the initial burn injury^7^, suggesting that the burn trauma itself may initiate a systemic effect that leads to increased risk of disease.

Evidence of a systemic response to burn injury has been reported using metabolic phenotyping platforms that characterise patterns of low-weight metabolites and metabolic intermediates in biofluids^8^. Such an approach is a useful tool for understanding the downstream pathophysiological changes associated with burn injury and can inform on physiological metabolic mechanisms in granular detail. Furthermore, the use of metabolic phenotyping platforms to detail the systemic response to burn injury may provide opportunity to develop metabolic biomarkers predictive of systemic burden and identify critical phases stages of systemic response, providing early identification of long-term complications and identifying novel pathway targets for therapeutic intervention^9^.

In particular, the use of metabolic phenotyping platforms have been previously used to highlight dysregulation of lipid metabolism that occurs following a burn injury^10^. This includes observations of increased levels of free fatty acids (FFAs) in the plasma hypothesised to result from triacylglyceride (TAG) breakdown in adipose and muscle^11, 12^, as well as acyl-carnitines in muscle tissue itself ^13^. In addition, accumulation of TAGs^13^ and lipoapoptotic-inducing palmitic and stearic lipids^14^ have been demonstrated to promote hepatic steatosis in the liver post-burn injury. Conversely a significant decline in FFAs, monoacylglycerides (MAGs), lysophosphatidylglycerols (LPGs) and lysophosphatidylethanolamines (LPEs) has been noted in burn-affected skin^15^.

Perturbations in circulating lipoproteins, macro-molecular lipid-protein complexes, have also been reported following burn injury. High-density lipoproteins (HDLs) were reported to be decreased during acute (total serum HDLs)^16^ and severe burn injuries^17^, whereas levels of very low-density lipoproteins (VLDL) and low-density lipoproteins (LDL) were found at higher concentrations in paediatric non-severe^18^ and adult severe burns^19^. The changes identified in lipids and lipoproteins are indicative of burn-induced hypocholesterolemia, hypophospholipidemia and hypertriglyceridemia, which have been associated with morbidity and mortality following a severe burn injury^20^.

However, most research into the metabolic consequences of burn injury has been focused towards severe burns^17, 21–28^, with a only small proportion examining changes in non-severe burns^16, 29, 30^. In addition, as the incidence of non-severe injury is much greater than severe, it is vital to understand how a less-severe injury impacts the systemic metabolic response, and the long-term health consequences of such injury events.

Therefore, to explore the systemic metabolic consequence of non-severe burn injury, we applied a multimodal strategy to compare the lipidic content of blood plasma from participants with a non-severe burn injury (classified as hospitalised with <15% total burn surface area) to non-burn controls. The approach used liquid chromatography-tandem mass spectrometry (LC-QQQ-MS) to characterise the plasma lipidome in combination with ^1^H nuclear magnetic resonance (NMR) spectroscopy to integrate lipoprotein and inflammatory signatures. Sampling of the non-severe burn injury group was performed following admission to hospital (2.89 days ± 2.75 days) and at a follow-up appointment (45.64 days ± 7.5 days), allowing for longitudinal evaluation of the systemic metabolic response to non-severe burn.

## Methods

### Patient recruitment

Participants were recruited from Western Australian State Adult Burns Unit at Fiona Stanley Hospital (Perth, Western Australia) as part of the placebo arm of the Celecoxib for Acute Burn Inflammation (CABIN) and fever study (Registration number: ACTRN12618000732280). Recruitment criteria included a burn injury clinically classed as non-severe (<15% TBSA), aged between 18 to 65 years, no significant co-morbidities (outlined in **Table S1**) and able to give consent.

The resultant non-severe burn cohort had a mean TBSA of 4.0% ± 3.6%. Baseline sampling occurred at 2.89 ± 2.75 days post-injury. Participants attended a follow-up clinical visit, 45.64 ± 7.5 days post initial admission, where a second sample was collected. The age at time of baseline collection was 39.87 ± 16.73 years. Non-burn controls (n=14) were recruited with no medical history of burn injury requiring hospital admission with the mean age matched to the burn cohort 40.09 ± 10.7 years. An overview of the study participant groups is presented in **Table 1**.

**Table 1.**
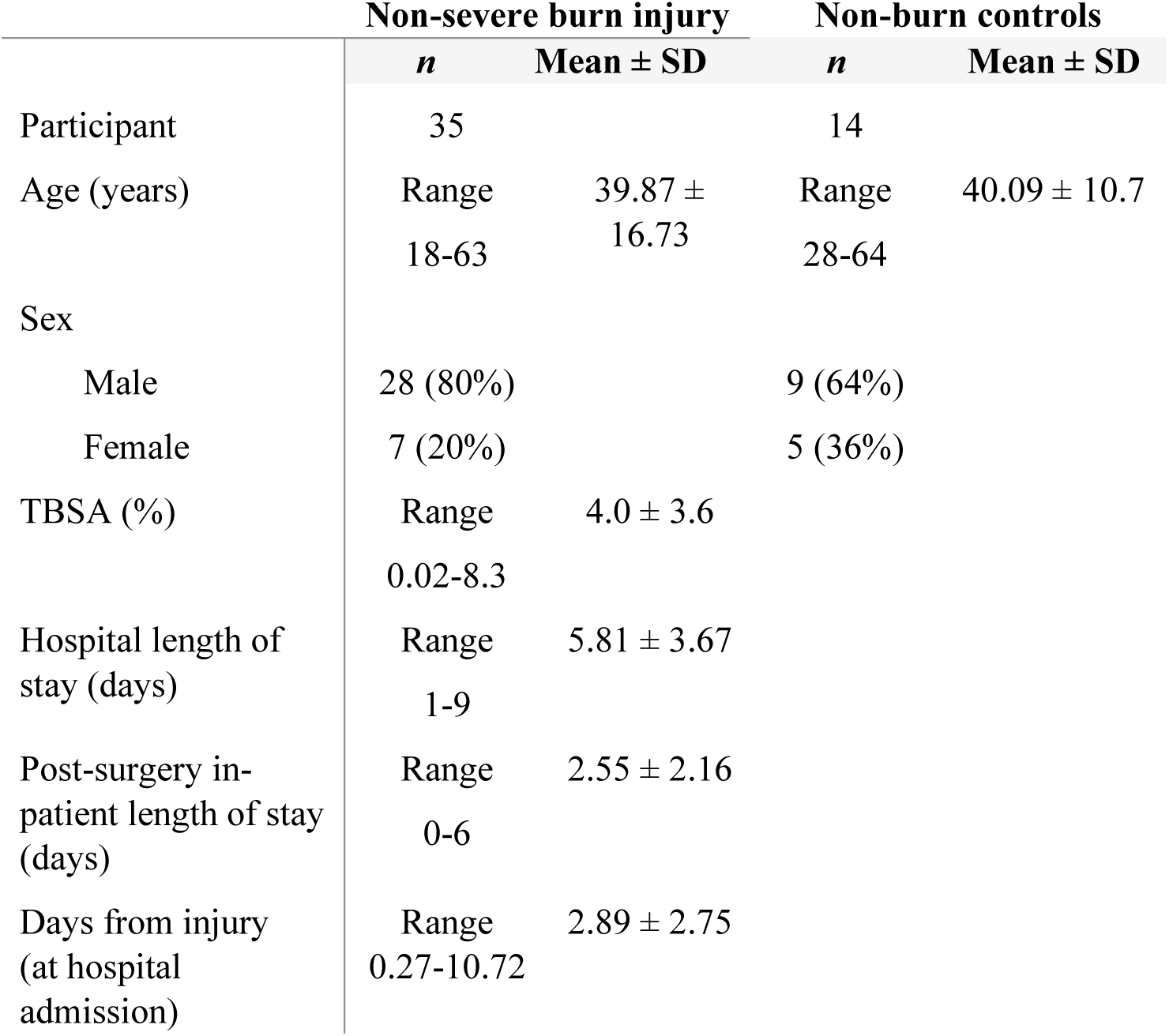
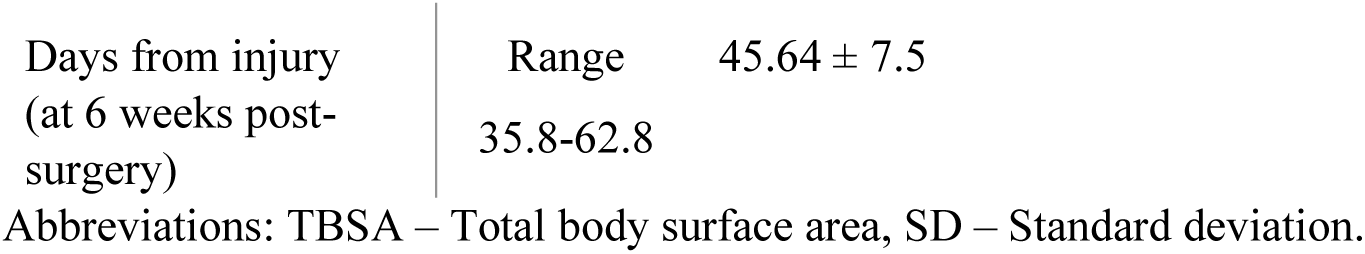
Demographics of non-severe burn patients from the CABIN Fever study and recruited non-burn controls.

Plasma samples were collected and prepared using a standardised protocol for both groups, whereby whole blood was collected into lithium heparin tubes, and centrifuged within 2 hours of collection. Plasma was aliquoted and stored at -80°C until analysis.

Ethics approval was obtained from South Metropolitan Health Service Health Research Ethics Committee (Approval number: RGS731) and Murdoch University Human Research Ethics Committee (Murdoch University Ethics no. 2020/053).

### Preparation of pooled quality control plasma samples

A commercial pooled human plasma (BioIVT, Westbury, NY, USA) and used as a quality control (QC) sample for analysis. Pooled QC was sub-aliquoted and underwent sample preparation for ^1^H NMR spectrometry or LC-QQQ-MS described below. QC samples were analysed at consistent intervals throughout the analytical run between every tenth study sample.

### 1H NMR spectroscopy of plasma

Plasma samples were prepared for ^1^H NMR spectroscopy analysis using a previously published method^31^. Briefly, analysis was performed on a 600 MHz Bruker Avance III HD spectrometer (Bruker BioSpin, Billerica, Massachusetts, USA) fitted with a BBI probe and a Bruker SampleJet™ robot (Bruker BioSpin, Billerica, Massachusetts, USA) using a cooling system maintaining samples at 5°C. Plasma samples were centrifuged at 13 000 x *g* for 10 min at 4°C and phosphate buffer (75 mM Na_2_HPO_4_, 2 mM NaN_3_, and 4.6 mM sodium trimethylsilyl propionate-[2,2,3,3-2H_4_] (TSP) in D_2_O, pH 7.4 ± 0.1) was added in a 1:1 ratio with the plasma supernatant. Of the mixture, 600 µL was then transferred into the Bruker SampleJet™ with SampleJet 5 mm outer diameter NMR tubes sealed with POM balls. NMR analysis was performed in accordance with the Bruker *in vitro* Diagnostics research (IVDr) methods, in which ^1^H 1D and Diffusion and Relaxation Editing (DIRE)^32–34^ experiments were run subsequently. ^1^H 1D experiments were run with solvent pre-saturation acquiring 32 scans, 98304 data points with 18028.85 Hz spectral width and experiment time 4 min 3 s^35^; and DIRE experiments acquired 64 scans into 98304 data points and 18028.846 Hz spectral width, experiment time 4 min 25 s^32–34^.

### Lipoprotein subfraction analysis

The ^1^H NMR spectroscopic analysis generates 112 lipoprotein parameters were generated for each plasma sample via the Bruker IVDr Lipoprotein Subclass Analysis (B.I.-LISA™), whereby the −CH_2_ at δ = 1.25 and −CH_3_ at δ = 0.80 peaks of the 1D spectrum after normalization to the Bruker QuantRef manager within Topspin, were quantified using a PLS-2 regression model^36^. The lipoprotein classes quantified were very low-density lipoproteins (VLDLs), low-density lipoproteins (LDLs), intermediate-density lipoproteins (ILDLs), and high-density lipoproteins (HDLs). VLDL subfractions were classified into six density classes (0.950 – 1.006 kg/L) of VLDL-1, VLDL-2, VLDL-3, VLDL-4, VLDL-5 and VLDL-6. LDL subfractions were assigned to six density classes of LDL-1 (1.019-1.031 kg/L), LDL-2 (1.031-1.034 kg/L), LDL-3 (1.034-1.037 kg/L), LDL-4 (1.037-1.040 kg/L), LDL-5 (1.040-1.044 kg/L), LDL-6 (1.044-1.063 kg/L). HDL subfractions were classified into four density classes of HDL-1 (1.063–1.100 kg/L), HDL-2 (1.100–1.125 kg/L), HDL-3 (1.125–1.175 kg/L), and HDL-4 (1.175–1.210 kg/L). An annotation list of lipoproteins from the B.I.-LISA™ method is provided in the Supplementary Information (**Table S2**). DIRE pre-processing included baseline correction using an asymmetric least-squares routine and the resulting spectra were normalised to the ERETIC signal using R (v4.4.1) with R Studio (v1.4.1)^37^. The GlycB acetyl signal (δ 2.07) arising from the glycoprotein acetyl residues were obtained from the DIRE experiment by integration. Due to overlap of triacylglyceride signals with GlycA (a composite peak of *N*-acetyl signals from five proteins: α-1-acid glycoprotein, α-1-antichymotrypsin, α-1-antitrypsin, haptoglobin, and transferrin), only the integrated signals from GlycB were used in all analyses. Supramolecular phospholipid composite (SPC) signal estimates were calculated from the peak spectral intensities in the regions of SPC_1_ δ 3.20–3.236, SPC_2_ δ 3.236–3.262, and SPC_3_ δ 3.262–3.30^38^.

### LC-QQQ-MS analysis of plasma lipids

Plasma lipids were analysed using a previously defined LC-QQQ-MS method for detection of 1163 lipid species^39^. Briefly, 10 µL of plasma was vortex mixed with 90 µL of propan-2-ol containing internal standards from Lipidyzer™ Internal Standard kit (SCIEX, MA, USA), SPLASH® LipidoMIX® (Sigma-Alridich, North Ryde, NSW, Australia) and Avanti Polar Lipids Lyso PG 17:1 and Lyso PS 17:1 (Sigma-Aldrich, North Ryde, NSW, Australia). Full description of internal standards is provided in **Table S3**. Samples were centrifugated at 14000 x *g* for 10 min and supernatant transferred to a 96-well plate. The LC-QQQ-MS method was run on a SCIEX ExionLC™ and QTRAP 6500+ using reversed-phase chromatographic separation and electrospray ionisation with polarity switching. Data were acquired using SCIEX Analyst® (v1.7.1).

### Data integration

For lipids analysed by LC-QQQ-MS, raw data were integrated in Skyline (v21.2)^40^ and concentrations of the detected lipids were calculated from the analyte to internal standard ratios. Data were pre-processed using R (v4.4.1) in R Studio (v1.4.1)^37^ where samples/features were filtered out if >50% missing values were present. Subsequently the remaining samples with <50% missing values were imputed by minimum value/2 and a final filtering out of features that had a >30% relative standard deviation (RSD) across the replicate quality control (QC) plasma samples. Signal drift was controlled for using the Random Forrest method from the *statTarget* package in the R software environment^41^.

### Statistical analysis

Principal component analysis (PCA) and supervised orthogonal projection to latent structures discriminant analysis (OPLS-DA) modelling were performed independently for lipids and lipoproteins using the R *metabom8* package (v0.2) from https://github.com/tkimhofer. OPLS-DA models were cross-validated using the Monte Carlo algorithm in the metabom8 package using a 2/3 model/prediction split. Visualisation plots and univariate statistics were generated using R (v4.4.1) in R Studio (v1.4.1). Variable importance in projection (VIP) scores from the OPLS-DA models were calculated using SIMCA® (v17, Sartorius, Goettingen, Germany).

## Results and discussion

### Data pre-processing and evaluation of quality control

Raw data signal deconvolution and pre-processing resulted in a panel of 112 lipoproteins and 852 lipid features. Observation of quality control (QC) pooled plasma samples clustering in scores plots of principal component analysis (PCA) performed independently for ^1^H NMR lipoprotein data and LC-MS lipidomic data, indicating robust and reproducible analytical performance throughout the acquisition run (**Figure S1A, S1B**).

### Plasma lipoprotein signatures indicate persistent inflammation after non-severe burn injury

OPLS-DA modelling comparing baseline non-severe burn and non-burn groups revealed a discriminant lipoprotein signature, which was consistent at the follow-up time point (**Figure 2**). To address the model performance with imbalanced group numbers, bootstrapping with 100 random iterations of a balanced model (10 *vs* 10) were performed, and the R^2^X and AUROC statistics were averaged (**Figure S2**). Furthermore, both the control *vs* baseline and control *vs* follow up data were used to build Receiver Operating Characteristic (ROC) and Precision-Recall (PR) curves to determine the accuracy and precision in balanced and imbalanced models^42, 43^. For control *vs* burns admission, the overall model (**Figure S2A**; R^2^X = 0.905; AUROC = 1) were compared with the mean scores for the bootstrapped iterations (**Figure S2B**; R^2^X = 0.68, AUROC = 1). For control *vs* burns follow up the overall model (**Figure S2C**; R^2^X = 0.854; AUROC = 1) were compared with the mean scores for the bootstrapped iterations (**Figure S2D**; R^2^X = 0.8; AUROC = 1). For both cases, the ROC-PR curves indicated the model using complete dataset is valid.

**Figure 1.**
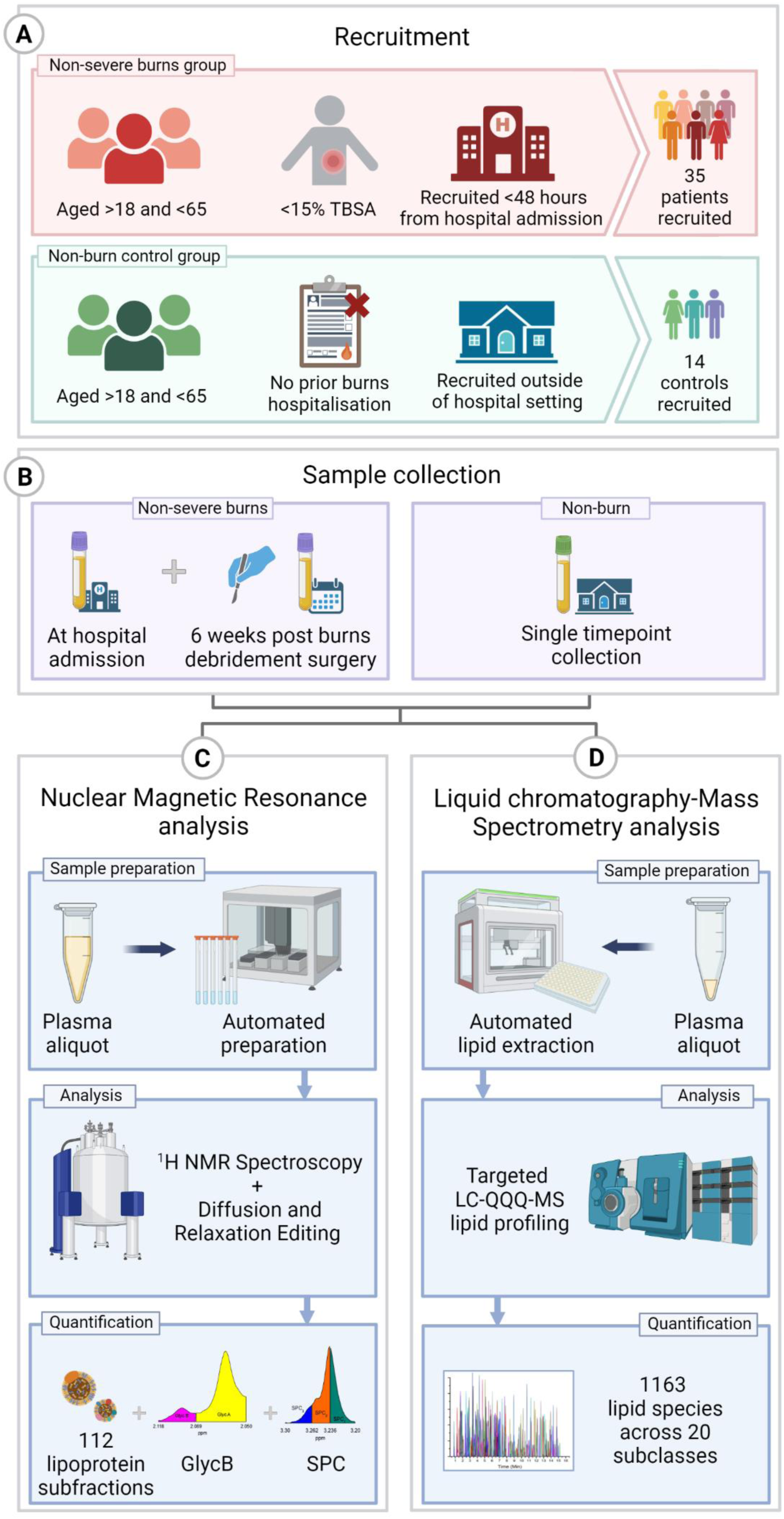
Overview of non-severe burn patient and non-burn participant recruitment and experimental workflow. **A**) Main selection criteria for non-severe burns and non-burn participants with final recruitment numbers used in the study. **B**) Sample collection timepoints for both groups with single plasma collection timepoint for non-burn controls and two timepoints from hospital admission and 6 weeks post burn debridement surgery for the non-severe burn group. **C**) Basic workflow of non-severe burn and non-burn plasma samples analysed by ^1^H nuclear magnetic resonance (NMR) spectroscopy and using Diffusion and Relaxation Editing. Data analysed included 112 lipoprotein subfractions, GlycB and the supramolecular phospholipid composite (SPC). **D**) Basic lipidomic workflow of non-severe burn and non-burn plasma analysed by liquid chromatography tandem mass spectrometry to quantify 1163 lipid species across 20 subclasses. Imaged created with BioRender.com

**Figure 2.**
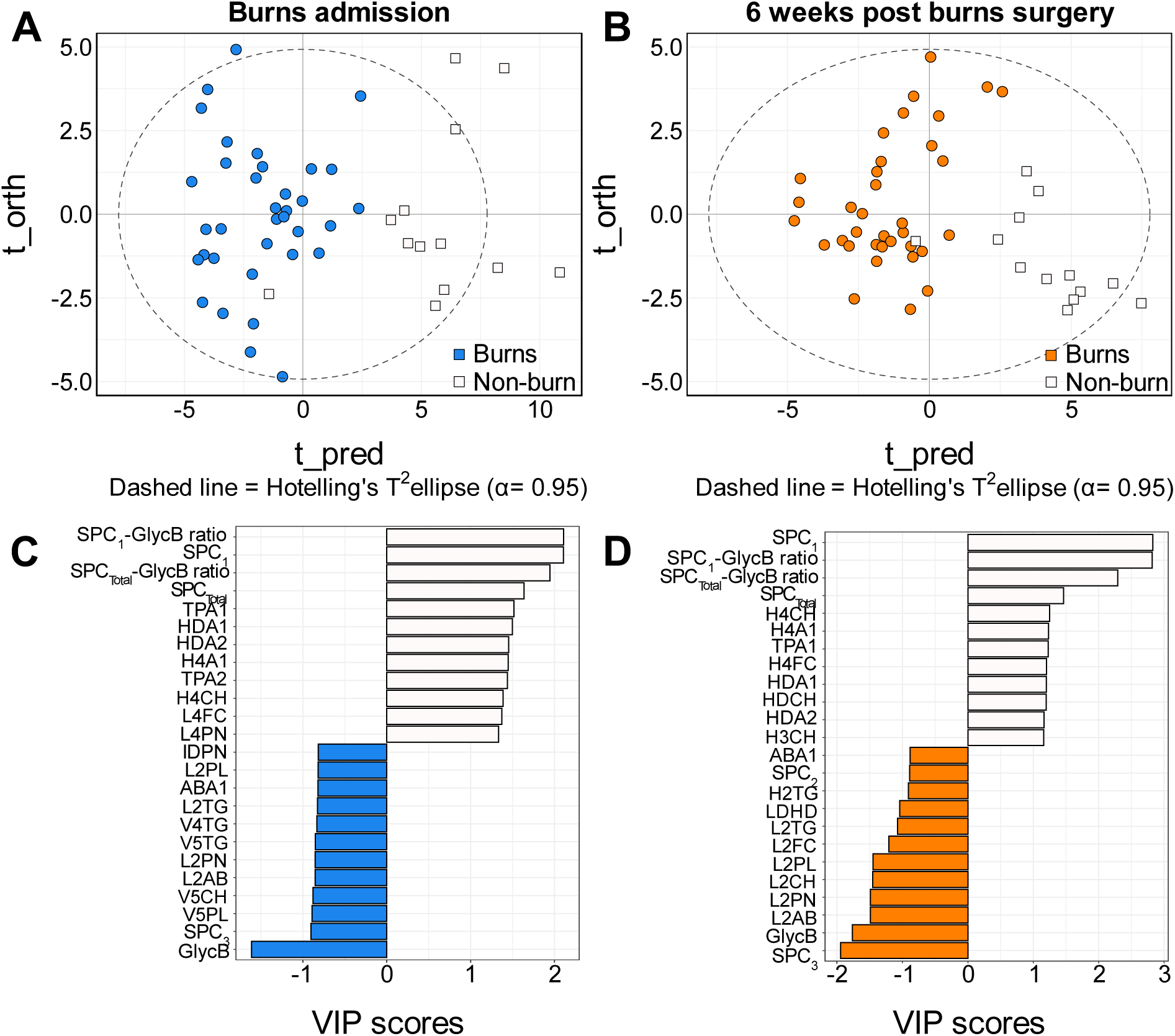
OPLS-DA analysis of lipoprotein profiles from paired burn plasma samples at admission to hospital and 6 weeks post-burn surgery compared to non-burn controls. **A)** OPLS-DA score plots of admission (n=35; coloured blue) (R^2^X = 0.25, Cross Validated-Area under the receiver operating characteristic (CV-AUROC) = 0.95 compared to non-burn controls (n=14; coloured white). **B)** OPLS-DA score plots of 6 weeks post-burn surgery (n=35; coloured orange) (R^2^X = 0.12, CV-AUROC = 0.96) compared to non-burn controls (n=14; coloured white). **C)** Bar plot of the top 12 variable importance in projection (VIP) scores, which summarises and ranks the influence of the individual lipoproteins on the OPLS-DA model, representing burns at admission (blue) and non-burn control (white). **D)** Bar plot of the top 12 VIP scores representing the 6 weeks post-burn surgery (orange) and non-burn controls (white).

From OPLS-DA modelling of non-severe *vs* non-burn cohorts, lipoproteins, GlycB and supramolecular phospholipid composite signals (SPCs)^44^ were ranked by their influence on the model using variable importance in projection (VIP) scores (**Figure 2C, D, Table S4**). The plasma profiles from the burn injury participant group were defined by higher concentrations of GlycB, SPC_3_ (related to LDLs), and LDL subfraction-2 particles, whereas higher concentrations of HDL cholesterol and HDL apolipoprotein A1 and A2 and lower levels of the phospholipid SPC_1_ (related to HDLs) were characteristic of the non-burn controls (**Figure 2C, D**).

GlycB was one of the strongest features defining the burn injury group at both admission and at 6 weeks post-surgery. This acute phase glycoprotein signal at δ 2.07 in the NMR spectra arises from the glycoprotein acetyl residues and has been shown to correlate with the clinical inflammatory marker high sensitivity C-reactive protein (hs-CRP)^44, 45^. GlycB has been shown to be associated with a range of acute and chronic inflammatory responses, such as vascular inflammatory conditions^45–48^, chronic heart disease^49^, prostate cancers^50^, rheumatoid arthritis^51–53^, SARS-CoV-2 infection^32^ and polycystic ovarian syndrome^47, 54^. A sustained increase in serum GlycB signals has been reported in a recent study of burn injury in paediatric patients where abnormal levels were found to persist 3 years after the initial burn injury and directly correlate with inflammatory cytokine levels, particularly interleukin-6, tumour necrosis factor-α and interferon-γ^18^.

Complimentary to GlycB, supramolecular phospholipid composite (SPC) signals deconvoluted from optimised Diffusion and Relaxation Editing (DIRE) NMR spectroscopy experiments ^32^ were reported to be perturbed in the non-severe burn group when compared with non-burn controls. SPC signals represent trimethylammonium residues in phospholipid components of lipoproteins, and results in three signal components: SPC_1_ signals representative of phospholipid content of HDL-4; SPC_2_ signals representative of phospholipids from HDL-1,2 and 3; and SPC_3_ representative of phospholipids from LDLs^38^. The signals have been previously used both independently and in ratio with GlycB as a sensitive markers for inflammation in infection^32, 55^, suggesting that such signals may be of value in predicting patient outcomes in burn injury. Our data reported significant decreases of SPC_1_ and SPC_1_/GlycB ratio in the non-severe burn cohort **(Figure 3)**, both at baseline and follow-up timepoints. For SPC_3_, significant elevations were observed only the follow up group, indicating a delayed impact of burn injury on the phospholipid content of LDL particles, whilst no significant changes were associated with SPC_2_ throughout the study.

**Figure 3.**
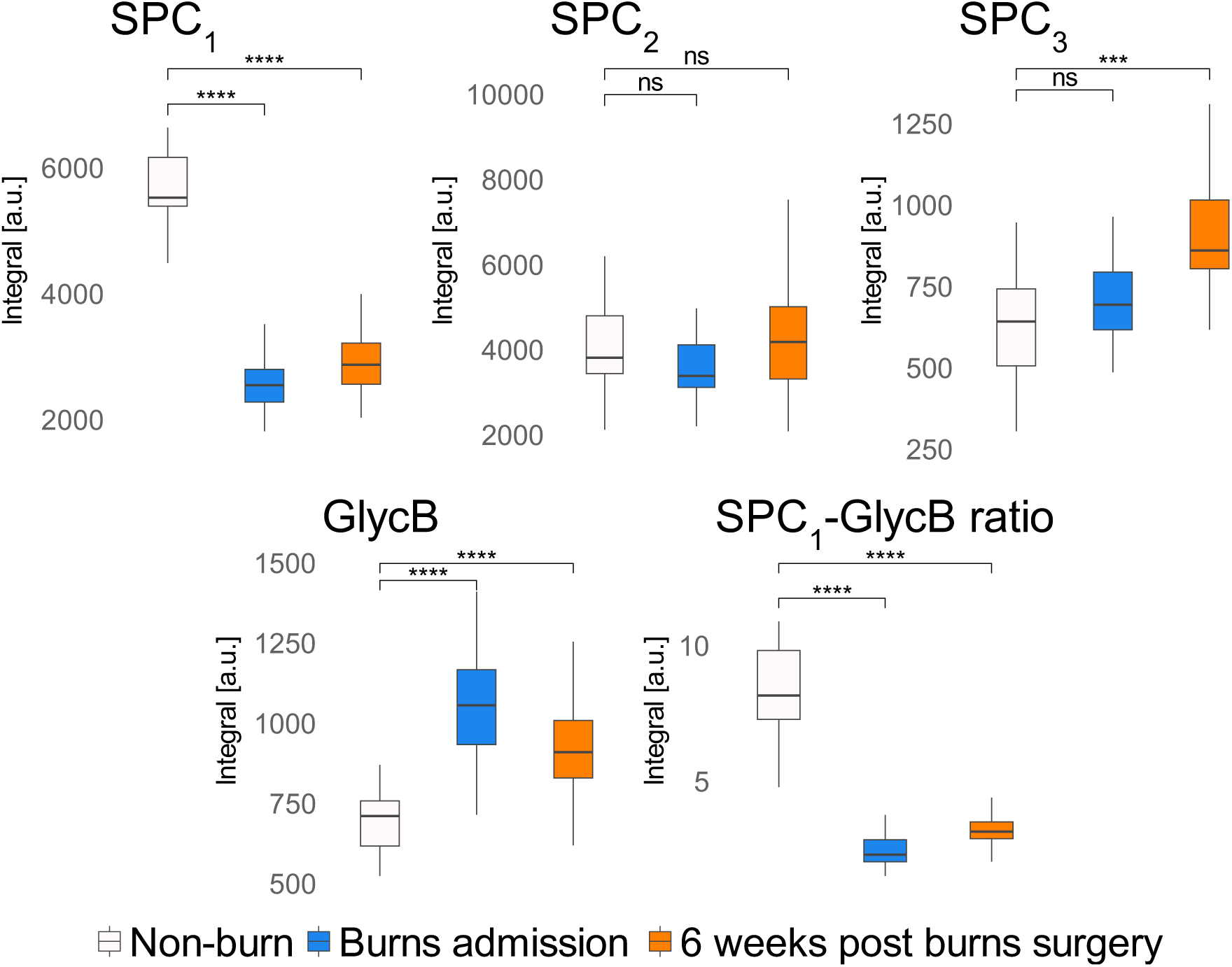
Univariate analysis of SPC_1_, SPC_2_, SPC_3_, GlycB and SPC_1_ to GlycB ratio values of non-burn controls compared to burns at admission and 6 weeks post-surgery. Box and whisker plots of non-burn controls (n = 14) (white) compared to burns at admission (n = 35) (blue) and 6 weeks post-surgery (n = 35) (orange) for SPC_1_, SPC_2_, SPC_3_, GlycB and SPC_1_ to GlycB ratio values. Significance level from Mann-Whitney U tests between non-burn controls to burns at admission and non-burn controls to 6 weeks post-surgery are shown above the corresponding plots and represented with ‘*’. Significance: ns = not significant; * = p-value < 0.05; ** = p-value < 0.01; *** = p-value < 0.001; **** = p-value < 0.0001.

Further discriminatory lipoprotein features in the NMR spectra included lower concentrations of HDL-associated apolipoprotein A1 (HDA1) and subfractions HDL-4 (HDL-4 apolipoprotein A1 (H4A1), HDL-4 free cholesterol (H4FC) and HDL-4 cholesterol (H4CH)) and higher concentrations of LDL-2 subfractions (LDL-2 phospholipids (L2PL), LDL-2 triacylglycerides (L2TG), LDL-2 particle number (L2PN), LDL-2 apolipoprotein B100 (L2AB), LDL-2 free cholesterol (L2FC) and LDL-2 cholesterol (L2CH)) in the non-severe burn group compared with non-burn. Key lipoprotein features were persistently perturbed when comparing both baseline admission and follow-up to non-burn controls indicating a sustained inflammatory signature (**Figure 2C, D**, **Figure 3**).

Such lipoprotein signatures have previously been associated with clinical outcomes in both acute and chronic inflammatory states, whilst HDL-associated apolipoprotein A1 has been reported to play an important anti-inflammatory role in diseased states, due to its ability to carry anti-inflammatory molecules, bioactive lipids and cholesterol^56, 57^. Depletions in plasma concentrations of specific subfractions including HDL-4 apolipoprotein A2 (H4A2) have also been associated with poor outcomes and mortality in patients with pulmonary arterial hypertension^58^, whilst lipoproteins H4A1, H4A2 and H4CH were significant markers of SARS-CoV-2 infection and severity ^32, 55, 59^. As with GlycB, our data indicted lower concentrations of HDL concentrations were sustained from admission to the follow-up sampling timepoint, indicating a persistence of the inflammatory response to non-severe burn injury.

In addition to acute inflammatory response, such lipoprotein subfraction signatures have also been associated with an increased risk of chronic disease. For example, LDL-2 subfractions (L2PL, L2TG, L2PN, L2AB, L2FC and L2CH) contain apolipoprotein B100, which is known to be atherogenic, and can be plaque-promoting and as such are linked with increased susceptibility to atherosclerosis^60, 61^. Our data revealed elevations of LDL particles in the burn injury group and highlights a potential avenue for enhanced atherosclerotic risk if an acute inflammatory state becomes persistent following a non-severe burn. In a similar theme, an inverse response between elevated LDL cholesterol and depleted HDL cholesterol has been previously reported as a risk marker of cardiovascular diseases^62^, such as ischemic heart disease^63^, myocardial infarction^62^ and coronary heart disease^64^. An increase in this LDL/HDL ratio (higher LDL and lower HDL cholesterol) is typically seen in chronic inflammatory conditions^65–67^, but has also been reported in survivors of severe burn injury (>20% TBSA) in which ratio levels related to worse healing outcomes, including longer length of hospital stay and higher infection rates^68^. Our analyses observed elevations in the LDL/HDL at both baseline and at follow-up, indicating a persistent inflammatory response that may have chronic implications for cardiovascular health in individuals who experience a non-severe burn injury.

This is further evidenced with evaluation of the apolipoprotein B100/apolipoprotein A1 (ABA1) ratio, which was elevated at both timepoints in our analyses (**Figure 2C**, **2D**). Apolipoprotein B100 (AB100) is a structural component to LDL that is reported to be atherogenic^69–71^, whilst apolipoprotein A1 is a component of HDL particles, and reported to be anti-atherogenic, anti-inflammatory and act as an antioxidant^70, 71^. Elevations in the ratio have been observed in both acute (SARS-CoV-2 infections^38^, severe burn injury^21, 22^), and chronic (metabolic syndrome^72^, cancers^73, 74^) inflammatory conditions. As with the LDL/HDL ratio, increases in the ABA1 ratio have also been proposed as a marker of abnormal lipoprotein changes in response to inflammation ^61, 70, 71, 75^ and risk of cardiovascular disease^76–78^. Such observations in our data may therefore be indicative of the potential metabolic avenues that influence the manifestation of the chronic inflammatory complications and cardiovascular complications linked to burn injury^4^.

### Plasma lipidomic signatures indicate persistent inflammation after non-severe burn injury

Our data indicated differences in lipidomic profiles between plasma samples collected from the non-burn controls and non-severe burn injury group at both baseline admission and at post-surgery follow-up (**Figure 4**). The imbalanced OPLS-DA models for lipids underwent the same ROC-PR curve analysis with a bootstrapped balanced model to assess performance. The burns admission *vs* control models for imbalanced (**Figure S3A**; R^2^X = 0.708; AUROC = 1) and balanced (**Figure S3B**; R^2^X = 0.67, AUROC = 1) were similar, as well as the 6 weeks post-surgery *vs* control imbalanced (**Figure S3C**; R^2^X = 0.705; AUROC = 1) and balanced (**Figure S3D**; R^2^X = 0.52, AUROC = 1) models. For both cases, the ROC-PR curves indicated the model using complete dataset is valid.

**Figure 4.**
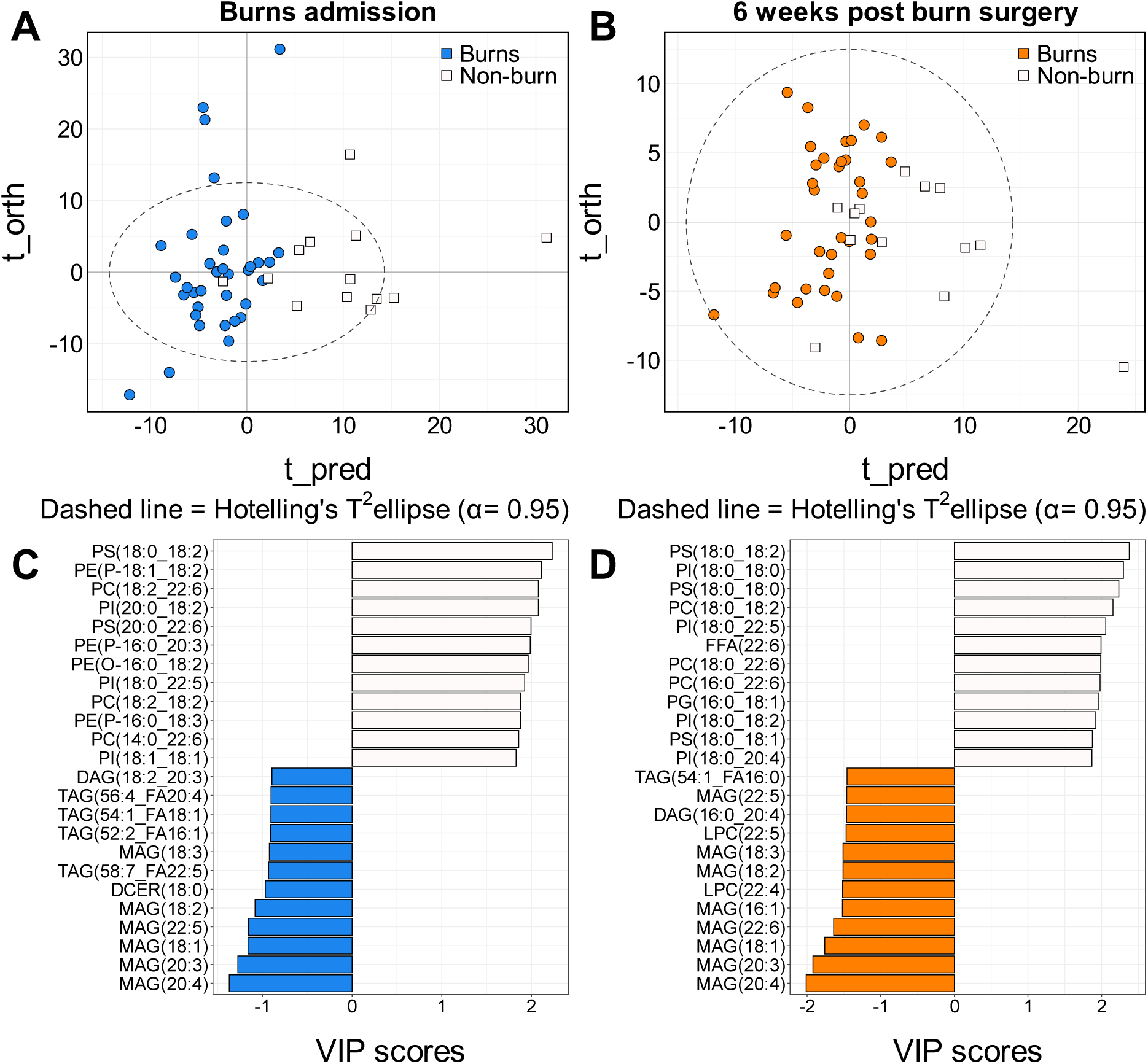
OPLS-DA analysis of lipids from plasma samples collected from hospitalised patients post-non-severe burn injury both at admission to hospital and 6 weeks post-surgery *vs* non-burn controls. **A)** OPLS-DA score plots of admissions (n=35; coloured blue) *vs* non-burn controls (n=14; coloured white) (R^2^X =0.21, CV-AUROC = 0.95). **B)** OPLS-DA score plots of 6 weeks post burn surgery (n=35; coloured orange) *vs* non-burn controls (n=14; coloured white) (R^2^X = 0.11, CV-AUROC = 0.80). **C)** Bar plots corresponding to the top 12 loading variables of each OPLS-DA model (ranked by variable importance in projection (VIP) score), colours represent burn injury group at admission (blue) *vs* non-burn controls (white). **D)** Bar plots corresponding to the top 12 loading variables of each OPLS-DA model (ranked by VIP score), colours represent burn injury group at six weeks (orange) *vs* non-burn controls (white).

Corresponding variable importance plot (VIP) revealed the top 12 lipids contributing to each of the pairwise models: non-burn controls *vs* non-severe burn - admission (**Figure 4C, Table S4**) and non-burn controls *vs* non-severe burn – follow-up (**Figure 4D, Table S4**). VIP visualisation plots structured by lipid class (x) and lipid sidechain length (y), coloured by the burn and non-burn cohorts, and the dot size correlating with the VIP contribution of each lipid variable loading for each pairwise model revealed a shift in lipidomic signature at follow-up compared with baseline admission, with specific lipid species elevating at the follow-up timepoint when comparing to the baseline non-burn controls. This included lipids from diacylglyceride (DAG), lysophosphatidylcholine (LPC), monoacylglyceride (MAG), phosphatidylcholine (PC), phosphatidylethanolamine (PE) and triacylglyercide (TAG) subclasses with side chain lengths of 16:0, 16:1 18:0, 18:1 and 20:4 (**Figure S4**).

Notably, the sidechain length of 20:4, or free fatty acid (FFA) 20:4 (FFA(20:4)) is commonly known as arachidonic acid, a precursor to numerous lipid mediators (eicosanoids) that are fundamental to inflammation and have been implicated in many inflammatory diseases^79^. During the inflammatory response, FFA(20:4) is metabolised into eicosanoid families, mostly in inflammatory cells, through four pathways which yield, but are not limited to, prostaglandins, isoprostanes, thromboxane and leukotrienes^80^. Metabolites within these families are predominately pro-inflammatory, known to enhance vasodilation, oedema formation, chemoattraction of inflammatory cells, and innate lymphocyte migration and activation^80^.

Further to this MAG(20:4) was a key variable in OPLS-DA modelling and was significantly higher in concentration in the burn cohort (OPLS-DA VIP score at admission = 0.95; Mann-Whitney U p-value <0.0001, **Figure 5**). Higher MAG(20:4) concentrations were also key in driving the OPLS-DA model comparing post-surgery follow-up (OPLS-DA VIP score = 1.32; Mann-Whitney U p-value < 0.0001) to the non-burn controls (**Figure 5**). MAG(20:4) is an endogenous lipid that can activate cannabinoid receptors which has importance as an inflammatory modulator^81, 82^, thought to modulate inflammation as a ligand to the pro-inflammatory cannabinoid 1 (CB1) receptor and the anti-inflammatory CB2 receptor ^83^ and has been reported as elevated in instances of inflammation following infection^84^.

**Figure 5.**
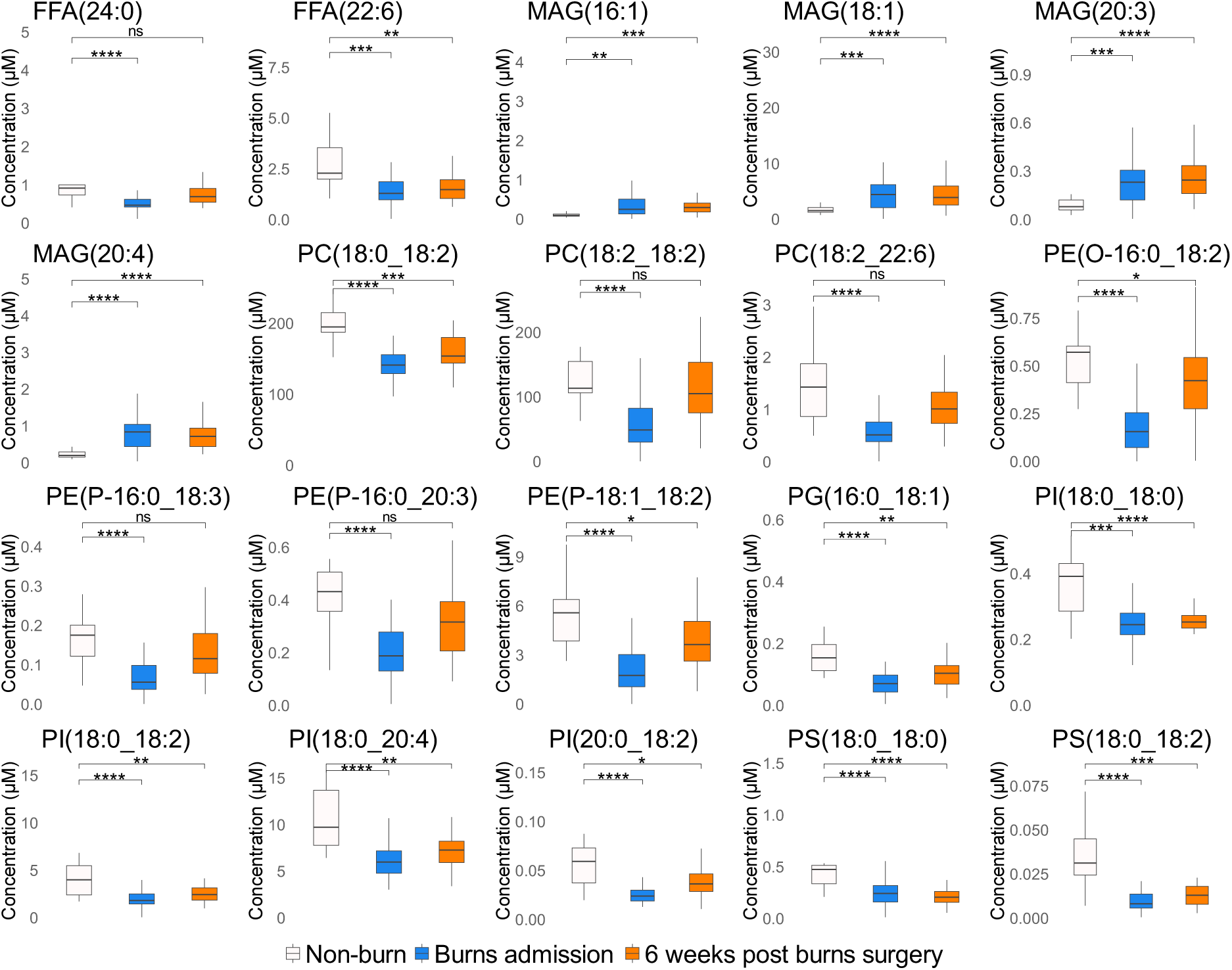
Top 20 lipids from univariate analysis of lipid profiles from non-burn controls compared to burns at admission and 6 weeks post-surgery. Box and whisker plots of non-burn controls (n = 14) (white) compared to burn admissions (n = 35) (blue) and 6 weeks post-burn surgery (n = 35) (orange) for the top 20 lipids from univariate analysis of non-burn controls *vs* the burn injury group at each timepoint. Significance level from Mann-Whitney U tests between non-burn controls to burn admissions and non-burn controls to 6 weeks post-burn surgery are shown above the corresponding plots and represented with ‘*’. Significance: ns = not significant; * = p-value < 0.05; ** = p-value < 0.01; *** = p-value < 0.001; **** = p-value < 0.0001.

As with the lipoproteins, evidence has accumulated of the pro-atherogenic impact that elevated levels of MAG(20:4) have *in vivo,* mainly through its activation of CB1^85–87^. It is hypothesised that MAG(20:4) stimulates monocytes and macrophages to migrate and promote pro-inflammatory cytokine expression, which can result in local vascular inflammation and atherosclerotic plaque formation^85, 88^. High concentrations of MAG(20:4) have been found in patients with different cardiovascular diseases, such as acute coronary syndrome^86^, obesity-induced cardiac dysfunction^89^ and chronic heart failure^90^, further indicating a possible avenue for the manifestation of chronic inflammatory and cardiovascular disease risk reported in individuals exposed to burn injury^4^.

In addition to MAG, TAG species were also reported to be higher in the non-severe burn group at baseline admission compared with the non-burn controls (**Figure 4C**). This complemented observations of higher concentrations of very low-density lipoprotein-5 (VLDL-5) subfractions (VLDL-5 cholesterol (V5CH) and VLDL-5 phospholipids (V5PL)) in the non-severe burn group at admission (**Figure 2C**). VLDLs consist mainly of TAGs (50-70% of the particle mass)^91^, with plasma VLDL remnants metabolised into IDL and then LDLs through the removal of triacylglycerides^92^. Elevated plasma TAGs and VLDL subfractions 1 and 2 have been correlated in non-burn inflammatory conditions, such as atherosclerotic cardiovascular disease, and has been hypothesised to be a result of impaired clearance by inefficient lipolysis^92^, again highlighting the association of common pathways that co-exist between acute injury and chronic inflammatory states.

Whilst specific acylglycerol species were higher in the non-severe burn injury groups, specific phospholipids were present at lower concentrations, including PC, PE, PG, PI and PS subclasses, and lyso-derivatives. This signature was maintained at both the admission and follow-up timepoints. PCs, PGs, PIs and PSs have all been reported to have anti-inflammatory properties^93–95^, but their influence on systemic inflammatory control and clinical outcomes post-burn injury have not been reported. Similar phospholipid signatures have been observed in infection, particularly in SARS-CoV-2 infections^84^, with PE(P-16:0_18:3), PE(P-18:1_18:2) and PC(18:2_18:2) as specific examples. The lower concentrations of phospholipid subclasses (PCs, PEs, PSs, PIs and PGs) observed is complimentary to observations of lower concentrations of HDL-4 subfractions observed (**Figure 2C**, **2D**), with plasma HDLs predominantly composed of phospholipids (40-60% of particle mass) and act as critical contributors regulating HDL cholesterol efflux ^96^.

In addition to PCs, their lyso-derivatives, LPCs, became one of the main drivers of separation between the burn and non-burn cohorts at the follow up timepoint (**Figure 4**). The biological role of the specific lipid species LPC(22:4) and LPC(22:5) has not been described in the literature to date, but the LPC family has been shown to be pro-inflammatory and potentially pro-atherogenic^97^. Promotion of atherogenesis has been associated with an increase in LPCs, through their ability to stimulate immune cell migration, particularly T-cell recruitment for apoptosis^97^. The change in signature from baseline to follow up may indicate secondary pathways activating post-non-severe burn injury, with implications towards persistent inflammation and shared pathways with chronic co-morbidities such as atherogenesis.

Finally, FFA(22:6) was significantly lower at both admission (Mann-Whitney U p-value < 0.001) and at the follow-up timepoint (Mann-Whitney U p-value < 0.01) compared to the non-burn controls (**Figure 4**, **5**). FFA(22:6) is also known as docosahexaenoic acid, a potent anti-inflammatory lipid mediator^80, 98^. This FFA can be metabolised into the eicosanoids resolvin and protectin, which can inhibit neutrophil and dendritic cell migration and modulate surface markers on lymphocytes to contribute to inflammatory resolution^80^, suggesting an ongoing persistent inflammatory response to injury, that is systemic even in cases of non-severe injury.

### Study limitations

We acknowledge that this study has limitations, particularly with respect to the bias in the sex of participants recruited to the study. The non-severe burn group was mostly comprised of males (80%). In Western Australia, adult males are hospitalised for burns at twice the rate of females, so this was expected in the non-severe burn cohort^1^. To counteract this, the non-burn control group was also weighted towards male participants (64.2%). Furthermore, to assess the influence of biological sex, each lipid and lipoprotein reported in the VIP analysis of OPLS-DA modelling were subjected to Mann-Whitney U testing (**Figure S5**), with no significant difference found between sex for each.

Furthermore, due to the acute nature of burn injury, intra-patient baseline prior to injury were not attainable. Additionally, there is risk that participants may have used non-steroidal anti-inflammatory medications for pain relief prior to hospital admission and enrolment. However, despite this, the primary signature observed indicated a systemic inflammatory response to injury that was persistent across the course of the study.

Finally, treatments for burn injury involve various interventions including surgery and implementation of a multi-modal analgesia treatment strategy. Conceptually, burn surgery may reduce the inflammatory stimulus by excising dead and damaged skin or conversely due to its invasive nature, surgery may provide a second inflammatory insult. It is therefore unclear whether increases of lipids and lipoproteins between admission and follow up are solely driven by burn injury or are influenced by the debridement surgery performed. However, the lipid and lipoprotein signatures were persistent throughout the study, indicating the primary driver of the signature was the injury. Further research will be required to determine the influence of surgery and the timing of surgery on metabolic phenotypes post-burn injury.

### Conclusions

Non-severe burn injury has both immediate and persistent influence on plasma lipid and lipoprotein phenotypes. Alterations in lipid metabolism indicate an inflammatory profile that is induced post-burn that is sustained at a systemic level. At the follow-up timepoint, the inflammatory marker GlycB remained significantly enhanced and corresponded with a decrease in the supramolecular phospholipid composite, specifically SPC_1_ and subsequently HDL-4 subfractions (H4A1, H4A2 and H4CH), that has previously been found as a distinguishing marker of systemic inflammation. The combination of low plasma HDL-4 subfractions, PC, PE, PG, PI, and PS concentrations, and high LDL-2 subfractions, MAGs (particularly MAG(20:4)) and LPCs following burn trauma may be suggestive of an increased risk of sustained inflammation leading to additional chronic co-morbities, including elevated incidence of pathways reported to be associated with atherosclerosis pathology and cardiovascular disease risk. This association may be mechanistic in the manifestation of chronic inflammatory and cardiovascular disease risk event reported in individuals exposed to burn injury in their lifecourse^4^, and warrants further epidemiological investigation. This potential risk and sustained inflammation underscores the value in monitoring lipid and lipoprotein levels following even non-severe burn injury. Such monitoring may provide opportunity to stratify patients by the systemic response to injury, and lead to personalised intervention strategies to improve acute care in cases of non-severe burn and enhance recovery outcomes.

## Supplementary Information

Table S1. Table of exclusion and inclusion criteria for recruitment of non-severe burn patients into the CABIN Fever study.

Table S2. Annotation of the keys used by the Bruker IVDr Lipoprotein Subclass Analysis (B.I.-LISA^TM^) method for each lipoprotein subclass.

Table S3. List of internal standards used for the LC-QQQ-MS method and associated part numbers.

Table S4. Combined variable importance in projection (VIP) scores of the 852 lipid species and 112 lipoproteins, SPCs and GlycB detected in both non-severe burns at admission and 6 weeks post-surgery, and non-burn cohorts after orthogonal projection to latent structures discriminant analysis (OPLS-DA).

Figure S1. Principal component analysis (PCA) of study samples and intermittently run quality control (QC) plasma samples to assess data robustness.

Figure S2. Model assessments of the original lipoprotein orthogonal projections to latent structures discriminate analysis (OPLS-DA) model with imbalanced burn and non-burn sample sizes and bootstrapped balanced burn and non-burn sample sizes.

Figure S3. Model assessments of the original lipid orthogonal projections to latent structures discriminate analysis (OPLS-DA) model with imbalanced burn and non-burn sample sizes and bootstrapped balanced burn and non-burn sample sizes.

Figure S4. Visualisation of the orthogonal projections to latent structures discriminate analysis (OPLS-DA) model variable important in projection (VIP) scores of each of the loading variables, plotted as lipid class (x-axis) vs lipid side chain (y-axis).

Figure S5. Box and whisker plots to assess the influence of sex on the most important lipoproteins, SPCs, GlycB and lipids using VIP scores and univariate statistics.

## Data Requirements

Raw data from the SCIEX Analyst® (v1.7.1) was converted to an open file format and uploaded to the Mass Spectrometry Interactive Virtual Environment (MassIVE) repository. Data identifiers are MSV000092486 and doi:10.25345/C5X34N282.

## Author Contributions

M.J. Ryan, E. Raby, G.L. Maker, F.M. Wood, E. Holmes, J.K. Nicholson, L. Whiley, M.W. Fear and N. Gray contributed to study concept and designed the research. E. Raby, M.W. Fear and F.M. Wood conceived, designed and led the CABIN Fever trial and provided the plasma samples and metadata. M.J. Ryan performed data mass spectrometry acquisition and drafted the manuscript. P. Nitschke and S. Lodge analysed the plasma samples using the NMR spectroscopy methods. M.J. Ryan and L. Whiley performed the pre-processing of MS data and statistical analysis of the lipid profiles. S. Lodge and R. Masuda performed the pre-processing of the NMR spectra. M.J. Ryan, S. Lodge, E. Holmes, J. Wist, J.K. Nicholson and R. Masuda performed modelling and interpretation of the lipoprotein subfractions, SPC and GlycB dataset. All authors edited and critically revised the manuscript. E. Raby, F.M. Wood, E. Holmes and J.K. Nicholson provided funding support for this manuscript.

## Acknowledgment

We would like to thank the Western Australian State Government and the Medical Research Future Fund for funding the work undertaken at the Australian National Phenome Centre. For the CABIN Fever trial, we thank the fund provided by the Department of Health Research Translation Programme, Fiona Wood Foundation and Spinnaker Health Research Foundation. We thank the Commonwealth of Australia for supporting MR with an Australian Government Research Training Program Scholarship. Lastly, we thank the Department of Jobs, Tourism, Science and Innovation, Government of Western Australian Premier’s Fellowship and Australian Research Council Laureate Fellowship funding for EH.

## Abbreviations

AB100: Apolipoprotein B100
ABA1: Apolipoprotein B100/apolipoprotein A1
B.I.-LISA: Bruker IVDr lipoprotein subclass analysis
CABIN: Randomised placebo-controlled trial of Celecoxib for acute burn inflammation
CER: Ceramide
CV-AUROC: Cross validated-area under the receiver operating characteristic
DAG: Diacylglyceride
DIRE: Diffusion and relaxation editing
FFA: Free fatty acid
H4A1: HDL-4 apolipoprotein A1
HDA1: HDL-associated apolipoprotein A1
HDL: High density lipoprotein
hs-CRP: high sensitivity C-reactive protein
ILDL: Intermediate low density lipoprotein
IVDr: *in vitro* Diagnostics research
L2AB: LDL-2-associated apolipoprotein B
LC-QQQ-MS: Liquid chromatography-tandem mass spectrometry
LDL: Low density lipoprotein
LPC: Lysophosphatidylcholine
LPE: Lysophosphatidylethanolamine
LPG: Lysophosphatidylglycerol
MAG: Monoacylglyceride
MS: Mass spectrometry
NMR: Nuclear magnetic resonance
OPLS-DA: Supervised orthogonal projection to latent structures discriminant analysis
PC: Phosphatidylcholine
PCA: Principal component analysis
PE: Phosphatidylethanolamine
PI: Phosphatidylinositol
PG: Phosphatidylglycerol
PS: Phosphatidylserine
QC: Quality control
RSD: Relative standard deviation
SM: Sphingomyelin
SPC: Supramolecular phospholipid composite
TAG: Triacylglyceride
TBSA: Total burn surface area
TSP: Sodium trimethylsilyl propionate-[2,2,3,3-2H_4_]
VIP: Variable important in projection
VLDL: Very low density lipoprotein

