## Supplementary Information Figure S1 - S5 and Table S1 - S3 for "Non-severe burns induce a prolonged systemic metabolic phenotype indicative of a persistent inflammatory response post-injury"

**Table of Contents**

**Supplementary Information** ........................................................................................................

Table S1. Table of exclusion and inclusion criteria for recruitment of non-severe burn patients into the CABIN Fever study........................................................................................................S-2

Table S2. Annotation of the keys used by the Bruker IVDr Lipoprotein Subclass Analysis (B.I.-LISA ™) method for each lipoprotein subclass. ..........................................................................S-2

Table S3. List of internal standards used for the LC-QQQ-MS method and associated part numbers. ........................................................................................................................S-5

Figure S1. Principal component analysis (PCA) of study samples and intermittently run quality control (QC) plasma samples to assess data robustness..............................................................S-7

Figure S2. Model assessments of the original lipoprotein orthogonal projections to latent structures discriminate analysis (OPLS-DA) model with imbalanced burn and non-burn sample sizes and bootstrapped balanced burn and non-burn sample sizes. .............................................................S-8

Figure S3. Model assessments of the original lipid orthogonal projections to latent structures discriminate analysis (OPLS-DA) model with imbalanced burn and non-burn sample sizes and bootstrapped balanced burn and non-burn sample sizes. .............................................................S-9

Figure S4. Visualisation of the orthogonal projections to latent structures discriminate analysis (OPLS-DA) model variable important in projection (VIP) scores of each of the loading variables, plotted as lipid class (x-axis) vs lipid side chain (y-axis). .........................................................S-10

Figure S5. Box and whisker plots to assess the influence of sex on the most important lipoproteins, SPCs, GlycB and lipids using VIP scores and univariate statistics. ............................................S-11

**Supplementary Information (excel)** ........................................................................................

Table S4. Combined variable importance in projection (VIP) scores of the 852 lipid species and 112 lipoproteins, SPCs and GlycB detected in both non-severe burns at admission and 6 weeks post-surgery, and non-burn cohorts after orthogonal projections to latent structures discriminate analysis (OPLS-DA) .........................................................................................................Tab 1

**Table S1. Table of exclusion and inclusion criteria for recruitment of non-severe burn patients into the CABIN Fever study.**

| Inclusion criteria | Exclusion criteria |
|---|---|
| <48 h from admission to hospital with burn injury | Pregnancy or breast feeding |
| 18 to 65 years of age | Renal dysfunction |
| Non-severe burn <15% TBSA | NSAID or sulphonamide allergy |
| Able to give consent | Current systemic corticosteroid use |
| | Suspected/confirmed haemophilia |
| | Current use of anticoagulant medicines |
| | Angina, myocardial infarction or heart failure |
| | History of gastric ulcer or gastrointestinal bleeding |
| | Regular NSAID use prior to injury |
| | Suspected/confirmed hepatic cirrhosis, portal hypertension or variceal bleeding |
| | Using angiotensin converting enzyme inhibitor or angiotensin II receptor blocker |

**Table S2. Annotation of the keys used by the Bruker IVDr Lipoprotein Subclass Analysis (B.I.-LISA™) method for each lipoprotein subclass.** Abbreviations: LDL – low-density lipoprotein; HDL – high-density lipoprotein; VLDL – very low-density lipoprotein; IDL – intermediate-density lipoprotein.

| Key | Class/Subclass | Compound | Concentration unit |
|---|---|---|---|
| TPTG | Total Plasma | Triglycerides | mg/dL |
| TPCH | Total Plasma | Cholesterol | mg/dL |
| LDCH | LDL | Cholesterol | mg/dL |
| HDCH | HDL | Cholesterol | mg/dL |
| TPA1 | Total Plasma | Apolipoprotein-A1 | mg/dL |
| TPA2 | Total Plasma | Apolipoprotein-A2 | mg/dL |
| TPAB | Total Plasma | Apolipoprotein-B100 | mg/dL |
| LDHD | Ratio LDL and HDL Cholesterol | LDL Cholesterol / HDL Cholesterol | -/- |
| ABA1 | Ratio of Apolipoproteins A1 and B100 | Apolipoprotein-A1 / Apolipoprotein-B100 | -/- |

| | | | |
|---|---|---|---|
| TBPN | Apolipoprotein-B100 carrying particles | Particle Number | nmol/L |
| VLPN | VLDL | Particle Number | nmol/L |
| IDPN | IDL | Particle Number | nmol/L |
| LDPN | LDL | Particle Number | nmol/L |
| L1PN | LDL-1 | Particle Number | nmol/L |
| L2PN | LDL-2 | Particle Number | nmol/L |
| L3PN | LDL-3 | Particle Number | nmol/L |
| L4PN | LDL-4 | Particle Number | nmol/L |
| L5PN | LDL-5 | Particle Number | nmol/L |
| L6PN | LDL-6 | Particle Number | nmol/L |
| VLTG | VLDL Class | Triglycerides | mg/dL |
| IDTG | IDL Class | Triglycerides | mg/dL |
| LDTG | LDL Class | Triglycerides | mg/dL |
| HDTG | HDL Class | Triglycerides | mg/dL |
| VLCH | VLDL Class | Cholesterol | mg/dL |
| IDCH | IDL Class | Cholesterol | mg/dL |
| LDCH | LDL Class | Cholesterol | mg/dL |
| HDCH | HDL Class | Cholesterol | mg/dL |
| VLFC | VLDL Class | Free Cholesterol | mg/dL |
| IDFC | IDL Class | Free Cholesterol | mg/dL |
| LDFC | LDL Class | Free Cholesterol | mg/dL |
| HDFC | HDL Class | Free Cholesterol | mg/dL |
| VLPL | VLDL Class | Phospholipids | mg/dL |
| IDPL | IDL Class | Phospholipids | mg/dL |
| LDPL | LDL Class | Phospholipids | mg/dL |
| HDPL | HDL Class | Phospholipids | mg/dL |
| HDA1 | HDL Class | Apolipoprotein-A1 | mg/dL |
| HDA2 | HDL Class | Apolipoprotein-A2 | mg/dL |
| VLAB | VLDL Class | Apolipoprotein-B100 | mg/dL |
| IDAB | IDL Class | Apolipoprotein-B100 | mg/dL |
| LDAB | LDL Class | Apolipoprotein-B100 | mg/dL |
| V1TG | VLDL-1 Subclass | Triglycerides | mg/dL |
| V2TG | VLDL-2 Subclass | Triglycerides | mg/dL |
| V3TG | VLDL-3 Subclass | Triglycerides | mg/dL |
| V4TG | VLDL-4 Subclass | Triglycerides | mg/dL |
| V5TG | VLDL-5 Subclass | Triglycerides | mg/dL |
| V1CH | VLDL-1 Subclass | Cholesterol | mg/dL |
| V2CH | VLDL-2 Subclass | Cholesterol | mg/dL |
| V3CH | VLDL-3 Subclass | Cholesterol | mg/dL |
| V4CH | VLDL-4 Subclass | Cholesterol | mg/dL |
| V5CH | VLDL-5 Subclass | Cholesterol | mg/dL |
| V1FC | VLDL-1 Subclass | Free Cholesterol | mg/dL |
| V2FC | VLDL-2 Subclass | Free Cholesterol | mg/dL |
| V3FC | VLDL-3 Subclass | Free Cholesterol | mg/dL |
| V4FC | VLDL-4 Subclass | Free Cholesterol | mg/dL |

| | | | |
|---|---|---|---|
| V5FC | VLDL-5 Subclass | Free Cholesterol | mg/dL |
| V1PL | VLDL-1 Subclass | Phospholipids | mg/dL |
| V2PL | VLDL-2 Subclass | Phospholipids | mg/dL |
| V3PL | VLDL-3 Subclass | Phospholipids | mg/dL |
| V4PL | VLDL-4 Subclass | Phospholipids | mg/dL |
| V5PL | VLDL-5 Subclass | Phospholipids | mg/dL |
| L1TG | LDL-1 Subclass | Triglycerides | mg/dL |
| L2TG | LDL-2 Subclass | Triglycerides | mg/dL |
| L3TG | LDL-3 Subclass | Triglycerides | mg/dL |
| L4TG | LDL-4 Subclass | Triglycerides | mg/dL |
| L5TG | LDL-5 Subclass | Triglycerides | mg/dL |
| L6TG | LDL-6 Subclass | Triglycerides | mg/dL |
| L1CH | LDL-1 Subclass | Cholesterol | mg/dL |
| L2CH | LDL-2 Subclass | Cholesterol | mg/dL |
| L3CH | LDL-3 Subclass | Cholesterol | mg/dL |
| L4CH | LDL-4 Subclass | Cholesterol | mg/dL |
| L5CH | LDL-5 Subclass | Cholesterol | mg/dL |
| L6CH | LDL-6 Subclass | Cholesterol | mg/dL |
| L1FC | LDL-1 Subclass | Free Cholesterol | mg/dL |
| L2FC | LDL-2 Subclass | Free Cholesterol | mg/dL |
| L3FC | LDL-3 Subclass | Free Cholesterol | mg/dL |
| L4FC | LDL-4 Subclass | Free Cholesterol | mg/dL |
| L5FC | LDL-5 Subclass | Free Cholesterol | mg/dL |
| L6FC | LDL-6 Subclass | Free Cholesterol | mg/dL |
| L1PL | LDL-1 Subclass | Phospholipids | mg/dL |
| L2PL | LDL-2 Subclass | Phospholipids | mg/dL |
| L3PL | LDL-3 Subclass | Phospholipids | mg/dL |
| L4PL | LDL-4 Subclass | Phospholipids | mg/dL |
| L5PL | LDL-5 Subclass | Phospholipids | mg/dL |
| L6PL | LDL-6 Subclass | Phospholipids | mg/dL |
| L1AB | LDL-1 Subclass | Apolipoprotein-B100 | mg/dL |
| L2AB | LDL-2 Subclass | Apolipoprotein-B100 | mg/dL |
| L3AB | LDL-3 Subclass | Apolipoprotein-B100 | mg/dL |
| L4AB | LDL-4 Subclass | Apolipoprotein-B100 | mg/dL |
| L5AB | LDL-5 Subclass | Apolipoprotein-B100 | mg/dL |
| L6AB | LDL-6 Subclass | Apolipoprotein-B100 | mg/dL |
| H1TG | HDL-1 Subclass | Triglycerides | mg/dL |
| H2TG | HDL-2 Subclass | Triglycerides | mg/dL |
| H3TG | HDL-3 Subclass | Triglycerides | mg/dL |
| H4TG | HDL-4 Subclass | Triglycerides | mg/dL |
| H1CH | HDL-1 Subclass | Cholesterol | mg/dL |
| H2CH | HDL-2 Subclass | Cholesterol | mg/dL |
| H3CH | HDL-3 Subclass | Cholesterol | mg/dL |
| H4CH | HDL-4 Subclass | Cholesterol | mg/dL |
| H1FC | HDL-1 Subclass | Free Cholesterol | mg/dL |
| H2FC | HDL-2 Subclass | Free Cholesterol | mg/dL |

| | | | |
|---|---|---|---|
| H3FC | HDL-3 Subclass | Free Cholesterol | mg/dL |
| H4FC | HDL-4 Subclass | Free Cholesterol | mg/dL |
| H1PL | HDL-1 Subclass | Phospholipids | mg/dL |
| H2PL | HDL-2 Subclass | Phospholipids | mg/dL |
| H3PL | HDL-3 Subclass | Phospholipids | mg/dL |
| H4PL | HDL-4 Subclass | Phospholipids | mg/dL |
| H1A1 | HDL-1 Subclass | Apolipoprotein-A1 | mg/dL |
| H2A1 | HDL-2 Subclass | Apolipoprotein-A1 | mg/dL |
| H3A1 | HDL-3 Subclass | Apolipoprotein-A1 | mg/dL |
| H4A1 | HDL-4 Subclass | Apolipoprotein-A1 | mg/dL |
| H1A2 | HDL-1 Subclass | Apolipoprotein-A2 | mg/dL |
| H2A2 | HDL-2 Subclass | Apolipoprotein-A2 | mg/dL |
| H3A2 | HDL-3 Subclass | Apolipoprotein-A2 | mg/dL |
| H4A2 | HDL-4 Subclass | Apolipoprotein-A2 | mg/dL |

**Table S2. List of internal standards used for the LC-QQQ-MS method and associated part numbers.**

| Internal standards | Part/Lot number |
|---|---|
| **Lipidyzer™ Internal Standard kit** | 5040156 |
| CE(16:0)-d7 | |
| CE(16:1)-d7 | |
| CE(18:1)-d7 | |
| CE(18:2)-d7 | |
| CE(20:3)-d7 | |
| CE(20:4)-d7 | |
| CE(20:5)-d7 | |
| CE(22:6)-d7 | |
| PC(16:0/16:1)-d9 | |
| PC(16:0/18:1)-d9 | |
| PC(16:0/18:2)-d9 | |
| PC(16:0/18:3)-d9 | |
| PC(16:0/20:3)-d9 | |
| PC(16:0/20:4)-d9 | |
| PC(16:0/20:5)-d9 | |
| PC(16:0/22:4)-d9 | |
| PC(16:0/22:5)-d9 | |
| PC(16:0/22:6)-d9 | |
| DAG(16:0/16:0)-d9 | |
| DAG(16:0/18:0)-d9 | |
| DAG(16:0/18:1)-d9 | |
| DAG(16:0/18:2)-d9 | |
| DAG(16:0/18:3)-d9 | |
| DAG(16:0/20:4)-d9 | |
| DAG(16:0/20:5)-d9 | |
| DAG(16:0/22:6)-d9 | |
| FFA(16:0)-d9 | |
| FFA(17:1)-d9 | |

| | |
|---|---|
| LPC(16:0)-d7 | |
| LPE(18:0)-d7 | |
| PE(18:0/18:1)-d7 | |
| PE(18:0/18:2)-d7 | |
| PE(18:0/18:3)-d7 | |
| PE(18:0/20:3)-d7 | |
| PE(18:0/20:4)-d7 | |
| PE(18:0/20:5)-d7 | |
| PE(18:0/22:5)-d7 | |
| PE(18:0/22:6)-d7 | |
| dSM(16:0)-d7 | |
| dSM(18:1)-d7 | |
| dSM(24:0)-d7 | |
| dSM(24:1)-d7 | |
| TAG(50:1-FA16:0)-d9 | |
| dTAG(52:1-FA18:0)-d9 | |
| TAG(52:2-FA18:1)-d9 | |
| TAG(52:3-FA18:2)-d9 | |
| TAG(52:4-FA18:3)-d9 | |
| TAG(54:4-FA20:3)-d9 | |
| TAG(54:(5-FA20:4)-d9 | |
| TAG(56:7-FA22:6)-d9 | |
| DCER(16:0)-d9 | |
| HCER(16:0)-d9 | |
| LCER(16:0)-d9 | |
| CER(d16:0)-d7 | |
| **SPLASH LipidoMIX** ™ internal standards | 330707 |
| PC(15:0-18:1)-d7 | |
| PE(15:0-18:1)-d7 | |
| PS(15:0-18:1)-d7 | |
| PG(15:0-18:1)-d7 | |
| PI(15:0-18:1)-d7 | |
| LPC(18:1)-d7) | |
| LPE(18:1)-d7 | |
| MAG(18:1)-d7 | |
| TAG(15:0-18:1)-d7-15:0 | |
| SM(d18:1-18:1)-d9 | |
| **Avanti Polar Lipids internal standards** | |
| LPS(17:1)-d7 | 858141C |
| LPG(17:1)-d7 | 858127C |

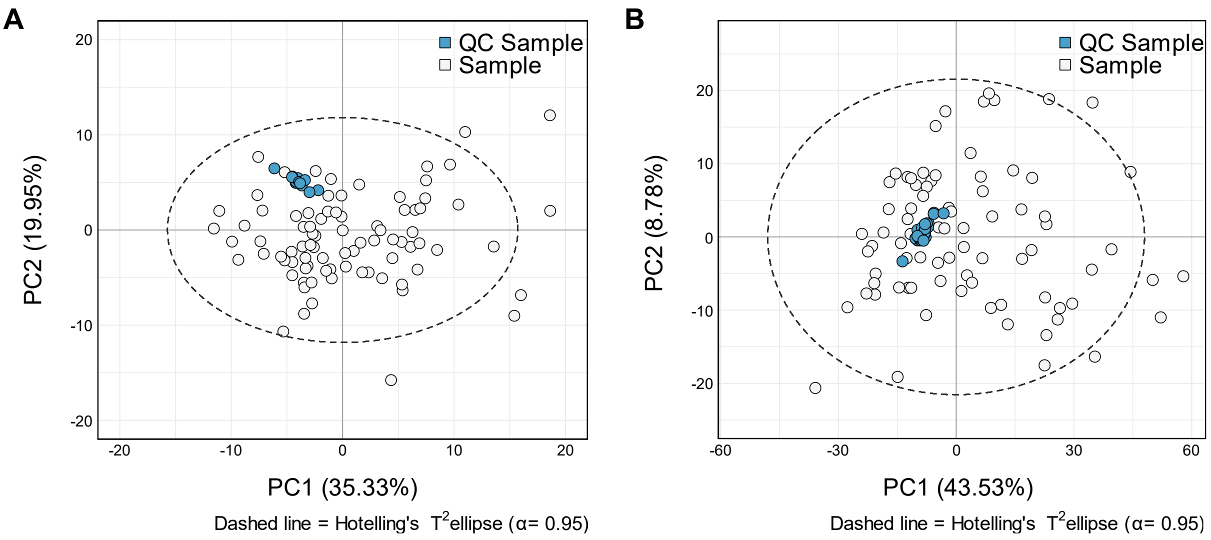


**Figure S1**. **Principal component analysis (PCA) of study samples and intermittently run quality control (QC) plasma samples to assess data robustness. A)** PCA scores plot of the lipoprotein profiles analysed by nuclear magnetic resonance (NMR) spectroscopy of intermittently run QC samples (coloured blue) and the study plasma samples (coloured white) (non-severe burns and non-burn cohorts). **B)** PCA scores plot of the lipid profiles analysed by liquid chromatography-tandem mass spectrometry (LC-QQQ-MS) of intermittently run QC samples (coloured blue) and the study plasma samples (coloured white) (non-severe burns and non-burn cohorts).

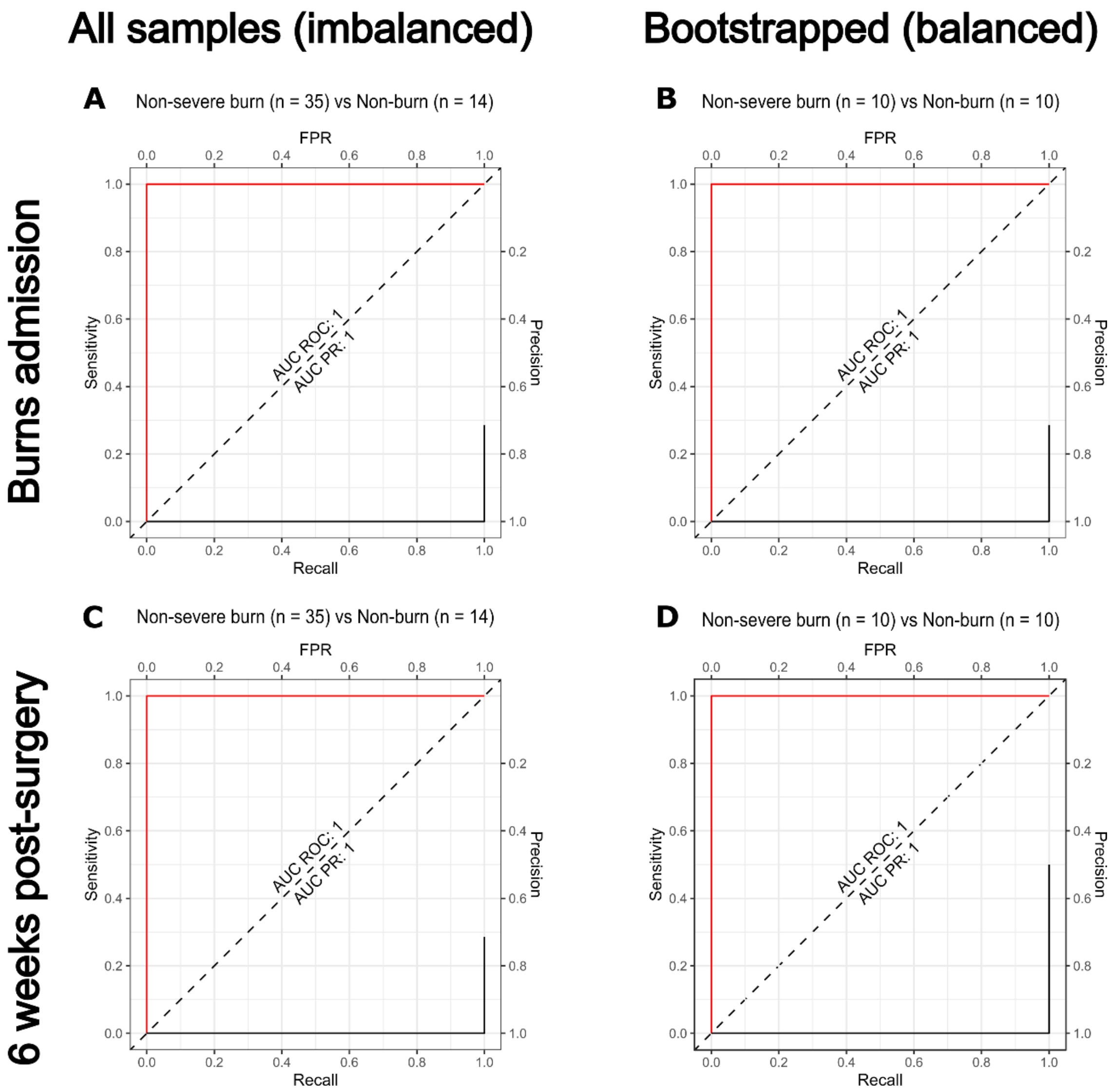


**Figure S2. Model assessments of the original lipoprotein orthogonal projections to latent structures discriminate analysis (OPLS-DA) model with imbalanced burn and non-burn sample sizes and bootstrapped balanced burn and non-burn sample sizes. A)** Receiver operating characteristic (Sensitivity vs False Positive Rate (FPR)) and Precision-Recall (ROC-PR) combined curve to assess accuracy in the OPLS-DA model separation for the original lipoprotein model (non-severe burns = 35; non-burn controls = 14) at burns admission ($R^2X = 0.905$; AUROC = 1). **B)** Using the lipoprotein profiles at burns admission, 100 iterations of the OPLS-DA model were bootstrapped with randomised samples in each cohort (non-severe burns = 10; non-burn controls = 10) in every iteration and analysed using the combined ROC-PR curve ($R^2X = 0.68$, AUROC = 1). **C)** ROC-PR combined curve to assess accuracy in the OPLS-DA model separation for the original lipoprotein model (non-severe burns = 35; non-burn controls = 14) at 6 weeks post-surgery ($R^2X = 0.854$; AUROC = 1). **D)** Using the lipoprotein profiles at 6 weeks post-surgery, 100 iterations of the OPLS-DA model were bootstrapped with randomised samples in each cohort (non-severe burns = 10; non-burn controls = 10) in every iteration and analysed using the combined ROC-PR curve ($R^2X = 0.8$, AUROC = 1).

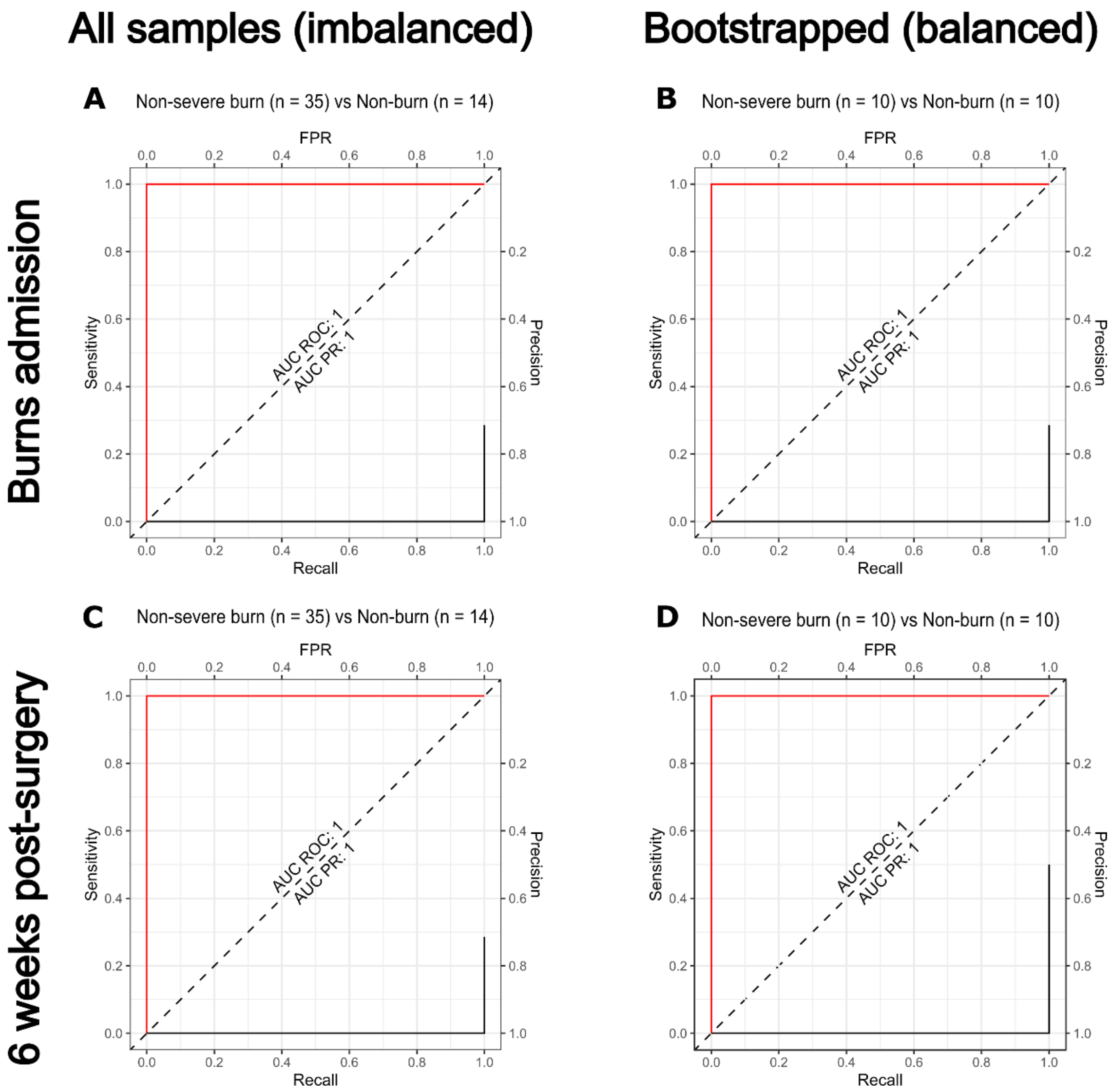


**Figure S3. Model assessments of the original lipid orthogonal projections to latent structures discriminate analysis (OPLS-DA) model with imbalanced burn and non-burn sample sizes and bootstrapped balanced burn and non-burn sample sizes. A)** Receiver operating characteristic (Sensitivity vs False Positive Rate (FPR)) and Precision-Recall (ROC-PR) combined curve to assess accuracy in the OPLS-DA model separation for the original lipid model (non-severe burns = 35; non-burn controls = 14) at burns admission ($R^2X = 0.708$; AUROC = 1). **B)** Using the lipid profiles at burns admission, 100 iterations of the OPLS-DA model were bootstrapped with randomised samples in each cohort (non-severe burns = 10; non-burn controls = 10) in every iteration and analysed using the combined ROC-PR curve ($R^2X = 0.67$, AUROC = 1). **C)** ROC-PR combined curve to assess accuracy in the OPLS-DA model separation for the original lipid model (non-severe burns = 35; non-burn controls = 14) at 6 weeks post-surgery ($R^2X = 0.705$; AUROC = 1). **D)** Using the lipid profiles at 6 weeks post-surgery, 100 iterations of the OPLS-DA model were bootstrapped with randomised samples in each cohort (non-severe burns = 10; non-burn controls = 10) in every iteration and analysed using the combined ROC-PR curve ($R^2X = 0.52$, AUROC = 1).

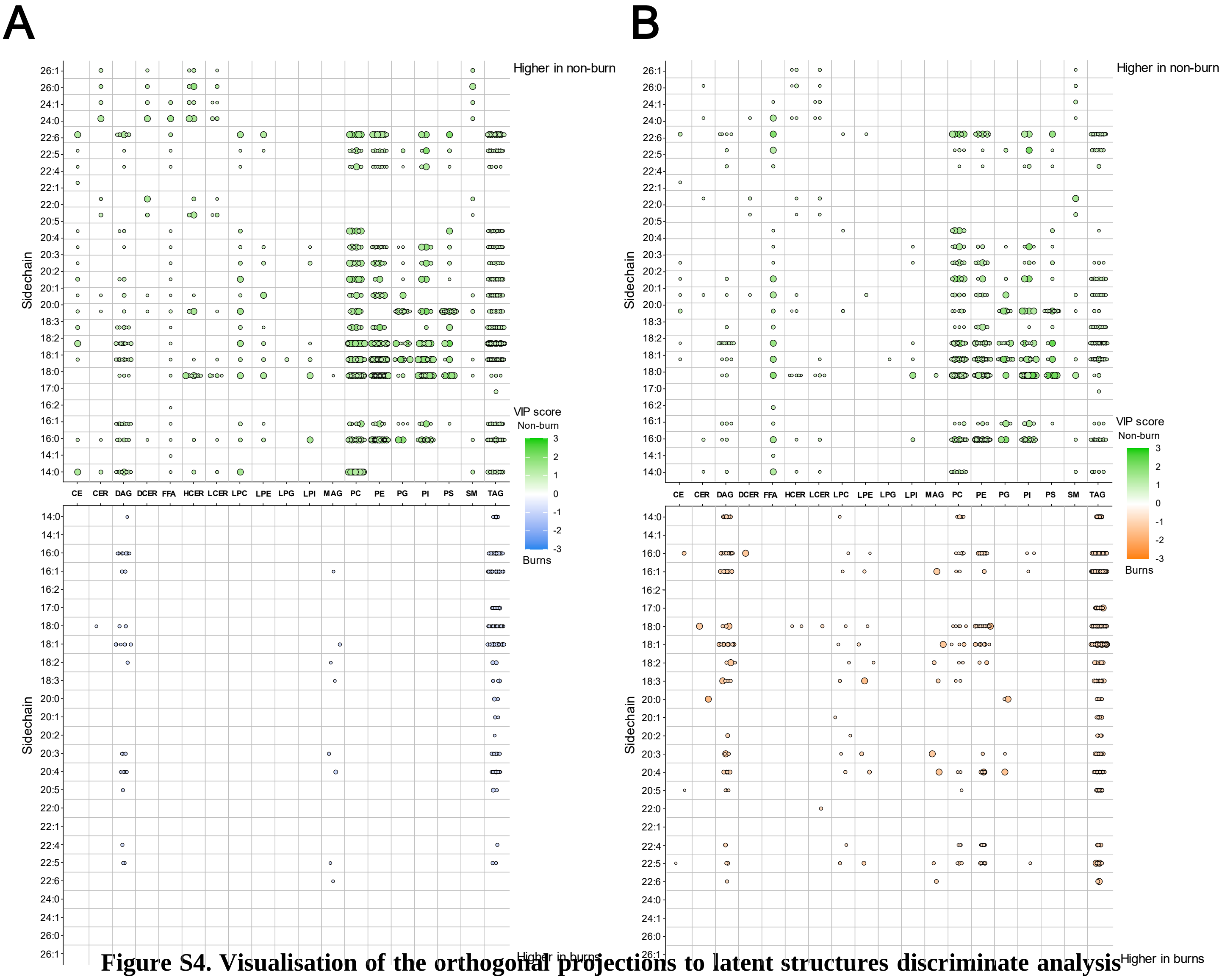


**Figure S4. Visualisation of the orthogonal projections to latent structures discriminate analysis (OPLS-DA) model variable important in projection (VIP) scores of each of the loading variables, plotted as lipid class (x-axis) vs lipid side chain (y-axis). A)** Top plot presents the VIP scores that drive the OPLS-DA direction of separation for the non-burn control group (green), whilst the bottom plot presents the VIP scores that drive the OPLS-DA direction of separation for burns admission (blue). **B)** Top plot presents the VIP scores that drive the OPLS-DA direction of separation for the non-burn control group (green), whilst the bottom plot presents the VIP scores that drive the OPLS-DA direction of separation for burn injury group at 6 weeks post-surgery (orange), with circle size and opacity proportional to the VIP score.

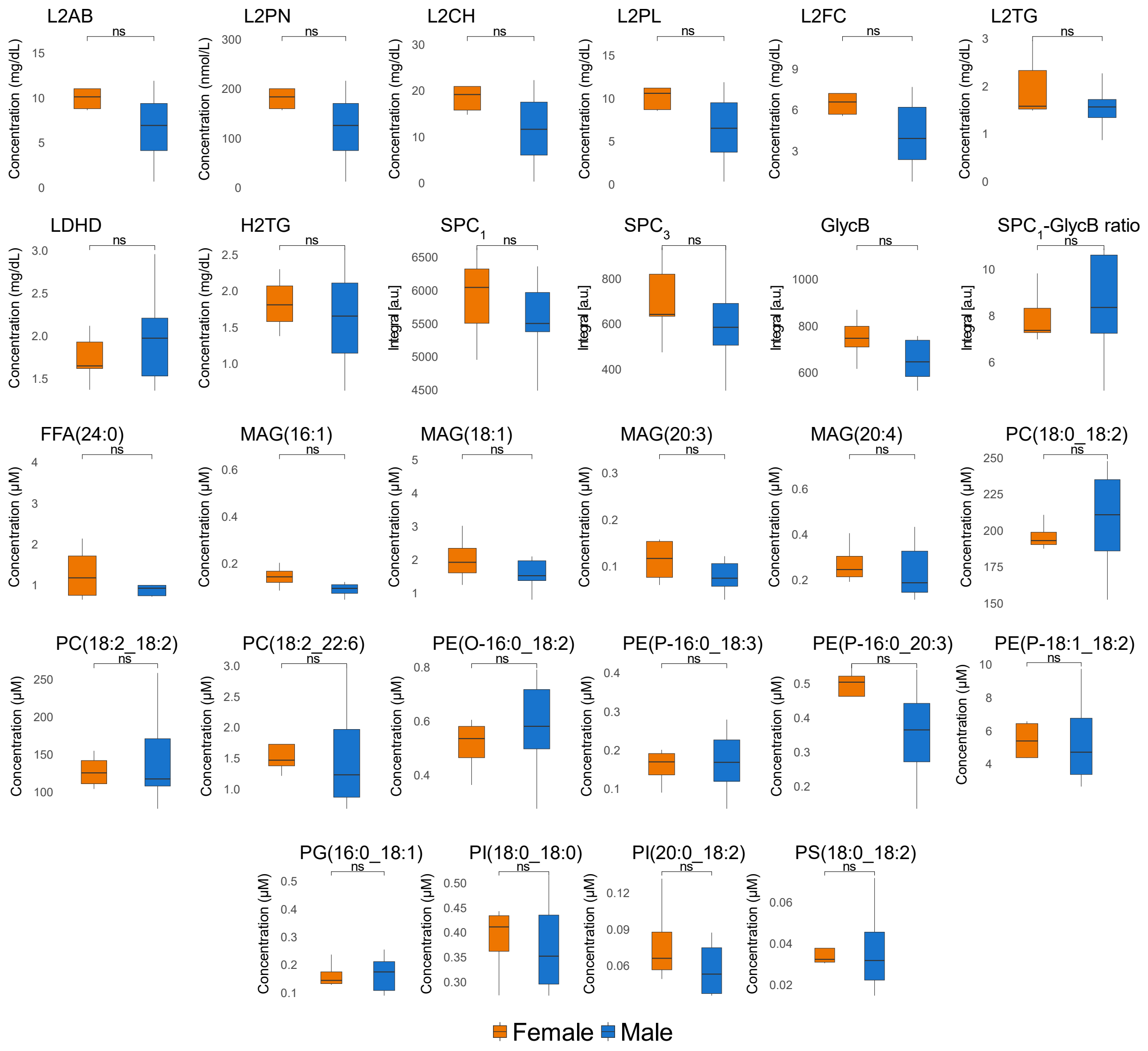


**Figure S5**. **Box and whisker plots to assess the influence of sex on the most important lipoproteins, SPCs, GlycB and lipids using VIP scores and univariate statistics**. Sex (F = 5 (orange); M = 9 (blue)) of non-burn controls were compared to determine any influence from sex using Mann-Whitney U. Significance level between sex are shown above the corresponding plots and represented with '*'. Significance: ns = not significant; * = p-value < 0.05; ** = p-value < 0.01; *** = p-value < 0.001; **** = p-value < 0.0001.
